# The minor and major spliceosome interact to regulate alternative splicing around minor introns

**DOI:** 10.1101/2020.05.18.101246

**Authors:** Anouk M. Olthof, Alisa K. White, Madisen F. Lee, Almahdi Chakroun, Alice K. Abdel Aleem, Justine Rousseau, Cinzia Magnani, Philippe M. Campeau, Rahul N. Kanadia

## Abstract

Mutations in minor spliceosome components are linked to diseases such as Roifman syndrome, Lowry-Wood syndrome, and early-onset cerebellar ataxia (EOCA). Here we report that besides increased minor intron retention, Roifman syndrome and EOCA can also be characterized by elevated alternative splicing (AS) around minor introns. Consistent with the idea that the assembly/activity of the minor spliceosome informs AS in minor intron-containing genes (MIGs), inhibition of all minor spliceosome snRNAs led to upregulated AS. Notably, alternatively spliced MIG isoforms were bound to polysomes in the U11-null dorsal telencephalon, which suggested that aberrant MIG protein expression could contribute to disease pathogenesis. In agreement, expression of an aberrant isoform of the MIG *Dctn3* by *in utero* electroporation, affected radial glial cell divisions. Finally, we show that AS around minor introns is executed by the major spliceosome and is regulated by U11-59K of the minor spliceosome, which forms exon-bridging interactions with proteins of the major spliceosome. Overall, we extend the exon-definition model to MIGs and postulate that disruptions of exon-bridging interactions might contribute to disease severity and pathogenesis.

## Introduction

The spliceosome is a ribonucleoprotein (RNP) complex responsible for the splicing of introns, a crucial step in the regulation of gene expression. Most eukaryotes contain two types of spliceosomes, that each recognize their own class of introns. The canonical spliceosome, also called the major spliceosome, splices major introns, whereas the minor spliceosome splices a small subset of introns called minor introns, that have divergent consensus sequences (Turunen et al, 2013). These minor introns are found in genes that are predominately made up of major introns, and thus the expression of these minor intron-containing genes (MIGs) requires the coordinated action of both the major and the minor spliceosome (Olthof et al, 2019). The splicing reactions executed by these two spliceosomes are analogous, as are the components. While the major spliceosome consists of the small nuclear RNPs (snRNPs) U1, U2, U4, U5 and U6, the minor spliceosome consists of the U11, U12, U4atac, U5 and U6atac snRNP (Tarn & Steitz, 1996a; Tarn & Steitz, 1996b). Most proteins are shared between the two spliceosomes, except the minor spliceosome contains seven unique proteins that are part of the U11/U12 di-snRNP (Will et al, 2004).

Even though the minor spliceosome is only responsible for splicing 0.5% of all introns in the human genome, its importance is underscored by several diseases linked to mutations in its components. For instance, germline mutations in the gene *RNU4ATAC*, which encodes the U4atac snRNA, have been linked to the diseases microcephalic osteodysplastic primordial dwarfism type 1 (MOPD1), Roifman syndrome and Lowry-Wood syndrome (Edery et al, 2011; Farach et al, 2018; He et al, 2011; Merico et al, 2015). Moreover, mutations in *RNU12* have been associated with an early-onset form of cerebellar ataxia (Elsaid et al, 2017). Finally, mutations in *RNPC3*, one of the minor spliceosome-specific proteins, have been linked to isolated growth hormone deficiency (IGHD) (Argente et al, 2014). While these diseases are the result of mutations in different genes, they all share a common molecular pathogenic mechanism, i.e. inhibition of the minor spliceosome. Transcriptomic analyses of peripheral blood mononuclear cells (PBMCs) from individuals with IGHD, cerebellar ataxia, MOPD1 and Roifman syndrome have revealed that this inhibition of the minor spliceosome results in elevated levels of retention of a subset of minor introns (Argente et al, 2014; Cologne et al, 2019; Elsaid et al, 2017; Merico et al, 2015). Despite a great degree of overlap in the MIGs that are affected in individuals with MOPD1 and Roifman syndrome, there is also a subset of MIGs that are only affected in one condition (Cologne et al, 2019). Thus, this might explain how inhibition of the same spliceosomal complex can result in unique symptoms and different levels of disease severity.

Alternatively, the type of mis-splicing, retention vs. alternative splicing (AS), that results from disruption of different minor spliceosome components might inform disease severity (Verma et al, 2018). For instance, mutations in *RNU12* and *RNPC3* are predicted to affect the first steps of spliceosomal assembly and intron recognition. This might not only result in intron retention, but also lead to AS, due to disruption of exon-definition interactions. In contrast, mutations in *RNU4ATAC* would affect the recruitment of the U4atac/U6atac.U5 tri-snRNP during later steps of the splicing reaction. Disruption of the tri-snRNP is not predicted to affect exon-definition interactions and therefore would solely result in intron retention. Indeed, transcriptomic analyses of individuals with MOPD1 and Roifman syndrome did show elevated minor intron retention and only minimal AS around minor introns (Cologne et al, 2019). In contrast, transcriptomic analysis of individuals with IGHD did not show the predicted increase in AS, but did show elevated minor intron retention (Argente et al, 2014). Thus, it remains unclear how disruption of different minor spliceosome components affects retention vs. AS outcomes in MIGs.

In order to specifically study retention and AS, we have previously developed a customized bioinformatics pipeline that was used to identify several tissue-specific AS events around minor introns (Olthof et al, 2019). Here, we leveraged this pipeline to understand the effect of minor spliceosome inhibition on minor intron retention and AS. We report that mutations in *RNU12*, besides leading to elevated minor intron retention, also resulted in the predicted increase in AS around minor introns. Unexpectedly, we also observed elevated AS around minor introns in an individual with Roifman syndrome. Finally, we explored the effect of *RNU4ATAC* mutations linked to Lowry-Wood syndrome, which had not previously been studied. We found that these mutations also resulted in elevated minor intron retention, but their effect on AS of minor introns was limited. These differences in molecular outcomes upon mutations in distinct minor spliceosome components were further corroborated in cell culture and *in vivo.* Specifically, we inhibited minor spliceosome components through antisense morpholinos and employed a conditional knockout (cKO) mouse for U11 snRNA. Here we report that alternatively spliced MIG transcripts were bound to polysomes in the U11-null dorsal telencephalon, suggesting they may be translated. Thus, we propose that the toxic gain-of-function of these novel MIG isoforms might contribute to the pathogenesis of minor spliceosome-related diseases. Finally, we uncovered that AS around minor introns is executed by the major spliceosome, whose activity can be regulated through the interaction with minor spliceosome component U11-59K. Together, these findings provide insight into the normal exon-definition interactions that take place to regulate the proper splicing of minor introns and the flanking major introns.

## Results

### The majority of retained minor introns in Lowry-Wood syndrome are also retained in Roifman syndrome

To determine the molecular consequences of mutation in *RNU4ATAC* on MIG expression and splicing of minor introns, we performed RNAseq on PBMCs from an individual with Roifman syndrome and an individual with Lowry-Wood syndrome, as well as their unaffected heterozygous parents and unrelated healthy controls. The individual with Lowry-Wood syndrome was a 28 year old male and a compound heterozygote for the *RNU4ATAC* variant n.120T>G and n.114G>C, whereas the individual with Roifman syndrome was 6 months old with two mutations *in trans* at n.17G>A and n.116A>G (Fig. S1A) (Magnani et al, 2009; Shelihan et al, 2018). While elevated levels of minor intron retention have previously been reported in individuals with Roifman syndrome, no transcriptomic analysis of individuals with Lowry-Wood syndrome has been reported so far (Merico et al, 2015). Therefore, we here compared the extent of minor intron retention in these individuals.

We detected retention of ∼170 minor introns in the healthy controls, Roifman syndrome carriers and Lowry-Wood syndrome carrier. This was drastically higher in the individuals with Roifman syndrome (319 minor introns) and Lowry-Wood syndrome (249 minor introns) (Fig. 1A). Moreover, the median level of minor intron retention (%MSI_Ret_) was significantly elevated in individuals with Roifman syndrome (57%) and Lowry-Wood syndrome (65%) compared to the carriers (25%, 20%, 33%) and unrelated healthy controls (22%, 20%, 22%) (*P*<0.0001) (Fig. 1B). In agreement with this finding, principal component analysis (PCA) of the %MSI_Ret_ values partitioned the patient samples from the healthy samples (PC1: 68% of the variance) (Fig. 1C). Thus, minor intron splicing is affected in both individuals with Roifman syndrome and Lowry-Wood syndrome.

**Figure 1.**
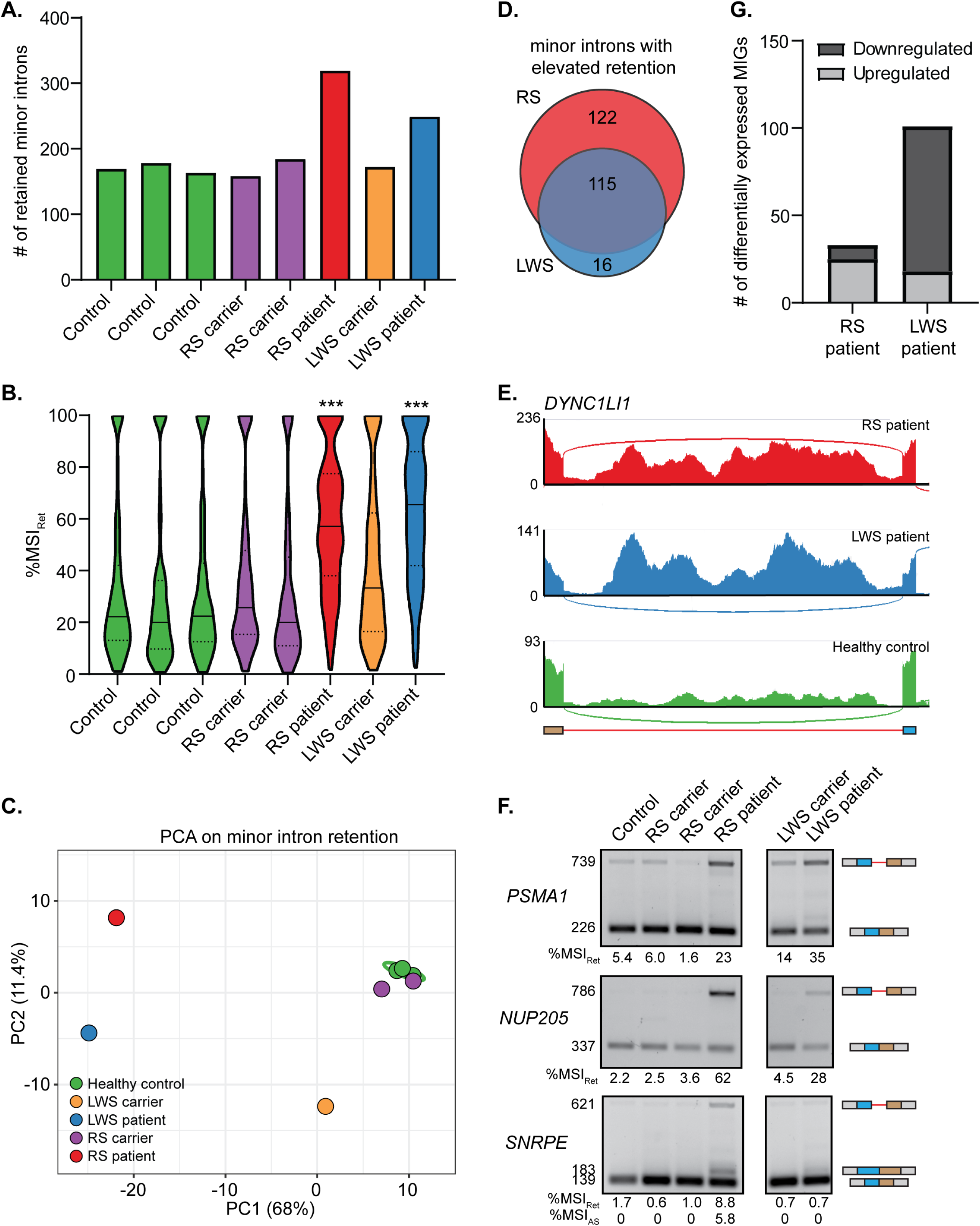
The majority of retained minor introns in Lowry-Wood syndrome are also retained in Roifman syndrome. **(A)** Bargraph with the number of retained minor introns in each individual. **(B)** Violin plots of the %MSI_Ret_ for all retained minor introns in each individual. Solid line denotes median; dotted lines depict Q1 and Q3. Statistical significance was determined by Kruskal-Wallis test, followed by post-hoc Dunn test. **\*\*\*=***P*<0.001. **(C)** Principal component analysis (PCA) on %MSI_Ret_ values in patients and controls. Prediction ellipse is drawn such that the probability is 95% that a new observation from the same condition falls inside the ellipse. **(D)** Venn diagram of the number of minor introns with elevated retention in the RS and LWS patient compared to healthy controls and the respective carriers. **(E)** Sashimi plots showing read coverage across minor introns. **(F)** Agarose gel images of RT-PCR products resulting from minor intron retention in PBMCs. Molecular weight of products is shown on the left; product schematics are shown on the right. The %MSI_Ret_ was calculated using ImageJ. **(G)** Bargraph with number of differentially expressed MIGs in the RS patient and LWS patient compared to healthy control. RS=Roifman syndrome; LWS=Lowry-Wood syndrome. See also Figure S1.

To determine which minor introns were affected by mutation in *RNU4ATAC*, we extracted those minor introns with a %MSI_Ret_ that was at least two-fold higher in the individuals with Roifman syndrome or Lowry-Wood syndrome compared to their respective carriers and the healthy controls. We detected 237 minor introns in 229 MIGs with elevated retention in the individual with Roifman syndrome, and 131 minor introns in 128 MIGs with elevated retention in the individual with Lowry-Wood syndrome (Fig. 1D; Dataset 1). Notably, 88% of all minor introns with elevated retention in the individual with Lowry-Wood syndrome were also retained at elevated levels in the individual with Roifman syndrome (Fig. 1D; Dataset 1). Conversely, 49% of all minor introns with increased minor intron retention in the individual with Roifman syndrome were also retained at elevated levels in the individual with Lowry-Wood syndrome (Fig. 1D; Dataset 1). This upregulation of minor intron retention was visualized for different MIGs by Sashimi plots and further validated by RT-PCR analysis (Fig. 1E-F, S1B). Since retention of introns often results in a premature stop codon, transcripts with retained introns are predicted to be degraded through pathways such as non-sense mediated decay (NMD) (Houseley & Tollervey, 2009). As such, we expected to observe downregulation of MIG transcripts with elevated minor intron retention. Surprisingly, the 229 MIGs that showed elevated minor intron retention in the individual with Roifman syndrome were not downregulated. Overall, only 8 MIGs were significantly downregulated in the individual with Roifman syndrome compared to healthy controls, which is consistent with previous reports where the minor spliceosome was inhibited (Fig. 1G; Dataset 2) (Baumgartner et al, 2018; Cologne et al, 2019; Markmiller et al, 2014; Merico et al, 2015). In contrast, 83 MIGs were significantly downregulated in the individual with Lowry-Wood syndrome compared to the healthy controls, although only 8 of these MIGs had evidence of minor intron retention (Fig. 1G; Dataset 2). Finally, PCA analysis on MIG expression did not separate the patients from the control on the first axis (Fig. S1C). While the second principal component did partition the two groups, it only explained 21% of the variance (Fig. S1B). Thus, minor intron retention is a better predictor of minor spliceosome disease than MIG expression.

### Alternative splicing around minor introns was elevated in Roifman syndrome and early-onset cerebellar ataxia, but not Lowry-Wood syndrome

Next, we wanted to evaluate other forms of AS around minor introns that might have been upregulated in response to minor spliceosome inhibition. Towards this goal, we employed our custom bioinformatics pipeline which also detects *de novo* AS events around minor introns. This revealed the presence of 66, 73 and 75 AS events around minor introns in healthy controls, and unaffected Roifman syndrome and Lowry-Wood syndrome carriers, respectively (Fig. 2A). This number was elevated in the individual with Roifman syndrome (137 AS events), but not in the individual with Lowry-Wood syndrome (78 AS events) (Fig. 2A). In total, we detected 188 AS events around minor introns in one or more samples, of which 79 were exclusively found in the disease state (Dataset 3). Specifically, 13 AS events were identified in both the individuals with Roifman syndrome and Lowry-Wood syndrome, whereas 14 and 52 AS events were exclusively found in the individual with Lowry-Wood syndrome and the individual with Roifman syndrome, respectively (Fig. S2A; Dataset 3). Moreover, 13 AS events were also detected in healthy controls or carriers, but the level of AS (%MSI_AS_) was upregulated at least 2-fold in the disease state (Dataset 3). Together, these 92 AS events affected the splicing of 68 minor introns through exon skipping (CAT2; 26%) or other AS types (Fig. S2B-C; Dataset 3). This suggested that multiple AS events could be employed in the same MIG. Indeed, cryptic 5’SS were frequently used in conjunction with cryptic 3’SS within the minor intron (Fig. 2B; Dataset 3). Furthermore, we confirmed these AS events common to both patients by RT-PCR, as well as those exclusively found in the individual with Roifman syndrome and not in the individual with Lowry-Wood syndrome (Fig. 2C). Regardless, PCA analysis on the %MSI_AS_ levels for all 188 detected AS events did separate the individuals with Roifman syndrome and Lowry-Wood syndrome from the controls on the first principal component (PC1: 45% of variance; Fig. S2D). Thus, even though the specific mutations in the U4atac snRNA may have a differential effect on the amount of AS around minor introns, it might contribute to disease pathogenesis.

**Figure 2.**
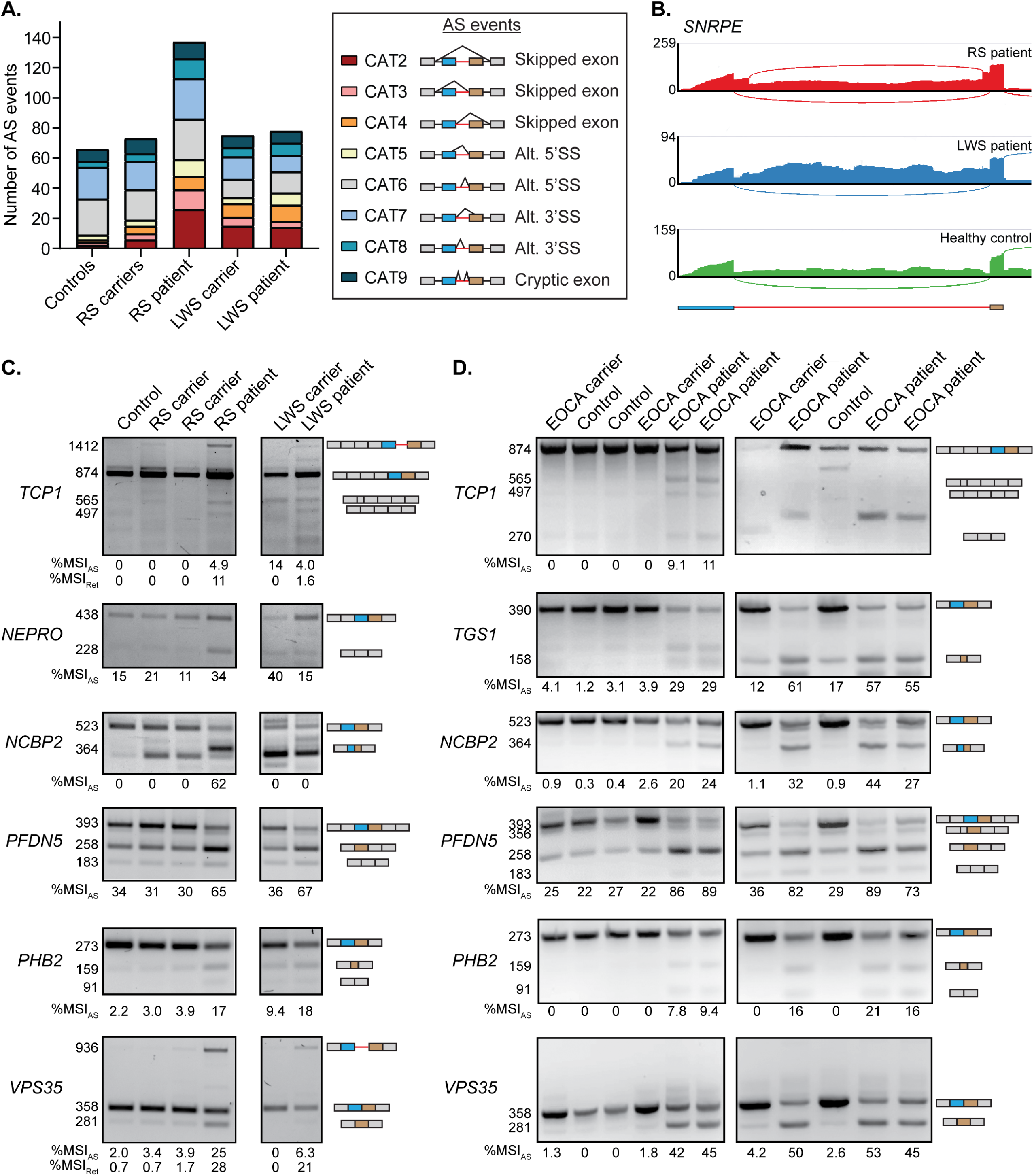
Alternative splicing around minor introns is elevated in Roifman syndrome and early-onset cerebellar ataxia, but not Lowry-Wood syndrome. **(A)** Bargraph with number of AS events around the minor intron in each individual. Schematics of different type of AS events are shown on the right. **(B)** Sashimi plots showing combined AS usage around minor introns. **(C-D)** Agarose gel images of RT-PCR products resulting from AS around minor introns in PBMCs of individuals with mutation in *RNU4ATAC* **(C)** and *RNU12* **(D)**. Molecular weight of products is shown on the left; product schematics are shown on the right. The %MSI_AS_ was calculated using ImageJ. RS=Roifman syndrome; LWS=Lowry-Wood syndrome; EOCA=early-onset cerebellar ataxia. See also Figure S2.

Finally, we wanted to determine whether minor spliceosome-related diseases caused by mutations in other minor spliceosome components than U4atac snRNA would also result in upregulated AS in MIGs. An autosomal recessive n.84C>T mutation in the U12 snRNA has been linked to early-onset cerebellar ataxia, and has been shown to lead to elevated minor intron retention in PBMCs of these patients (Fig. S2E) (Elsaid et al, 2017). However, the effect of this mutation on AS around minor introns had not yet been explored. To this end, for those minor introns that were alternatively spliced in the *RNU4ATAC* RNAseq data, we performed RT-PCR analysis on PBMCs from individuals with early-onset cerebellar ataxia, as well as their heterozygous parents and unrelated healthy controls. (Fig. 2A, S2A, S2D). We found that all of the AS events detected in the individual with Roifman syndrome, were also identified in individuals with early-onset cerebellar ataxia (Fig. 2C-D). In addition, a novel isoform for *TCP1* was detected in individuals with early-onset cerebellar ataxia from one family (Fig. 2D). Comparison of the %MSI_AS_ values for the events around the minor introns of *TCP1, NCBP2, PFDN5* and *VPS35* revealed that AS were generally higher in individuals with early-onset cerebellar ataxia than the individuals with Roifman syndrome and Lowry-Wood syndrome (Fig. 2C-D). Moreover, the minor intron retention events in *TCP1* and *VPS35* that we could detect in PBMCs from individuals with Roifman syndrome and Lowry-Wood syndrome, were absent in individuals with early-onset cerebellar ataxia (Fig. 2C-D). In all, these findings suggest that the mode by which the minor spliceosome is inhibited influences both the type of mis-splicing (retention vs. exon skipping), as well as the level of mis-splicing.

### Inhibition of minor spliceosome snRNAs each result in elevated AS around minor introns

To further explore the differential effect of minor spliceosome inhibition on the AS and retention levels of minor introns, we next employed antisense morpholino oligonucleotides (MO) against U12, U4atac and U6atac snRNAs in HEK293 cells. This was followed by capture of the nascent RNA and RNAseq (Fig. S3A). While this strategy would reduce the number of MIGs for analysis, it allowed us to capture the most immediate splicing defects after minor spliceosome inhibition. Compared to the control MO, we found that inhibition of U12 snRNA led to significantly elevated retention of 22 minor introns, whereas inhibition of U4atac and U6atac snRNA both resulted in the significant retention of 74 minor introns (Fig. S3B-D; Dataset 4). Importantly, an MO against U2 snRNA, a major spliceosome component, did not result in elevated minor intron retention (Fig. S3E).

Next, we analyzed the level of AS across minor introns in the aforementioned conditions. We found 94 AS events around minor introns in the control condition (Fig. 3A). Moreover, we detected 147, 119 and 152 AS events upon inhibition of U12 snRNA, U4atac snRNA, and U6atac snRNA, respectively (Fig. 3A). In contrast, inhibition of U2 snRNA resulted in a reduction of AS around minor introns (69 AS events) (Fig. 3A). The increased AS levels upon inhibition of the minor spliceosome components was primarily due to an increase in exon skipping (CAT2-CAT4) (Fig. 2A, 3A; Dataset 5). While also present in the control condition, the usage of this type of AS event had doubled upon minor spliceosome inhibition (Fig. 3A). Overall, 111 AS events were uniquely present when one or more minor spliceosomal components were inhibited (Dataset 5). In addition, 12 AS events were also present in the control condition, but significantly upregulated upon inhibition of the minor spliceosome (Dataset 5). Together, 65% of these events involved the skipping of one or both exons flanking the minor intron (Fig. S3F). To determine whether the individual minor spliceosome snRNAs regulated AS across different minor introns, we next performed an intersection analysis of all AS events that were upregulated upon inhibition of the minor spliceosome. This revealed that only 18% of the 123 upregulated AS events was observed in the U12, U4atac and U6atac MO condition, thereby affecting the splicing of 18 minor introns (Fig. 3B; Dataset 5). Instead, AS across the majority of minor introns was upregulated in just one or two MO conditions (Fig. 3B; Dataset 5). RT-PCR analysis revealed that these AS events were detected in all MO conditions, but at different levels, such that they did not always pass the stringent filtering criteria by RNAseq (Fig. 3C). For example, AS around the minor intron of *TGS1* was most highly upregulated upon inhibition of U4atac snRNA, whereas AS around the minor intron of *E2F3* was most responsive to inhibition of U12 snRNA (Fig. 3C). Thus, AS across minor introns is differentially affected by U12, U4atac and U6atac snRNA in a gene-specific manner.

**Figure 3.**
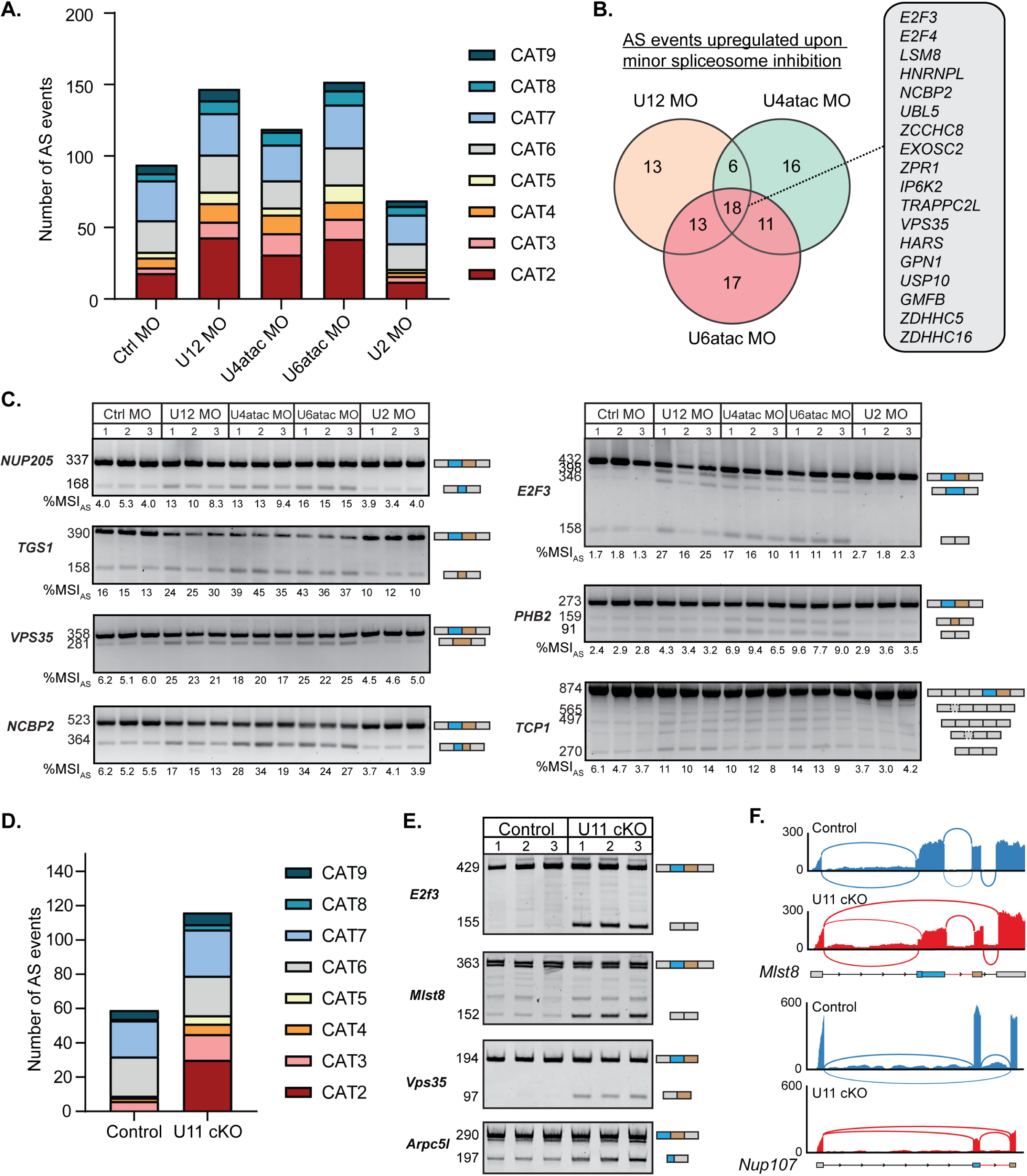
Inhibition of minor spliceosome snRNAs each result in elevated AS around minor introns. **(A)** Bargraph with number of AS events around the minor introns. AS categories are the same as in Fig. 2A. **(B)** Venn diagram showing the overlap of minor introns with significant upregulation of AS upon inhibition of the minor spliceosome. **(C)** Agarose gel images of RT-PCR products resulting from AS around minor introns. Molecular weight of Sanger-sequenced products is shown on the left; product schematics are shown on the right. The %MSI_AS_ was calculated using ImageJ. **(D)** Bargraph with number of AS events around the minor intron in U11 cKO embryos. **(E)** Acrylamide gel images of RT-PCR products resulting from AS around minor introns. Molecular weight of Sanger-sequenced products is shown on the left; product schematics are shown on the right. **(F)** Sashimi plots showing combined AS usage around minor introns. MO=morpholino. See also Figures S3 and S4.

Since MOs have not been proven effective against U11 snRNA, we instead analyzed RNAseq from the dorsal telencephalon of E12 U11 cKO embryos to study the effect of U11 inhibition on AS of minor introns (Baumgartner et al, 2018; Tarn & Steitz, 1996b). We found that U11 loss resulted in skipping of both exons flanking 30 minor introns (CAT2), while this was never observed in the control (Fig. 3D). In total, 116 AS events were observed in the U11 cKO, of which 50% were exclusive to the mutant and many were validated by RT-PCR and Sanger sequencing (Fig. 3E-F, Fig. S4A-B; Dataset 6). The remainder 58 AS events were detected in both samples (Fig. 3D). Of these, only three AS events, found in the MIGs *Mapk9, Spc24*, and *Ccdc43*, were significantly upregulated in the U11 cKO (Dataset 6). While it is hard to compare the specific AS events observed in the U11-null dorsal telencephalon to those observed in cell culture or in individuals with mutations in minor spliceosome components, these data do underscore that inhibition of all minor spliceosome snRNAs results in the production of aberrant MIG transcripts, especially through elevated levels of exon skipping.

### Alternatively spliced MIG transcripts are exported to the cytoplasm and bound to polysomes

Aberrant transcripts are normally quickly detected by quality control mechanisms, resulting in their degradation by the nuclear exosome or non-sense mediated decay (NMD) (Houseley & Tollervey, 2009). To test whether alternatively spliced MIG transcripts would be subject to exosome-mediated degradation in the nucleus, we fractionated the dorsal telencephalons from control and U11 cKO E12 embryos into a nuclear (NE) and cytoplasmic (CE) fraction (Fig. 4A). We found that all of the AS events detected in whole-cell extract were also detected in the CE of U11 cKO embryos (Fig. 3E, S4B, 4A). The successful export of these alternatively spliced MIG transcripts led us to investigate the effect of the AS on the open-reading frame (ORF), as frameshifts and premature stop codons are predicted to activate NMD (Chang et al, 2007). We found that ∼50% of the AS events were in frame, whereas the other half resulted in a frameshift (Fig. 4B). The majority of the AS events that caused a frameshift also introduced a premature stop codon that was predicted to activate NMD (Fig. 4C). In five MIGs, the minor intron was positioned close enough to the 3’ end of the transcript that the premature stop codon was not predicted to trigger the NMD pathway (Fig. 4C).

**Figure 4.**
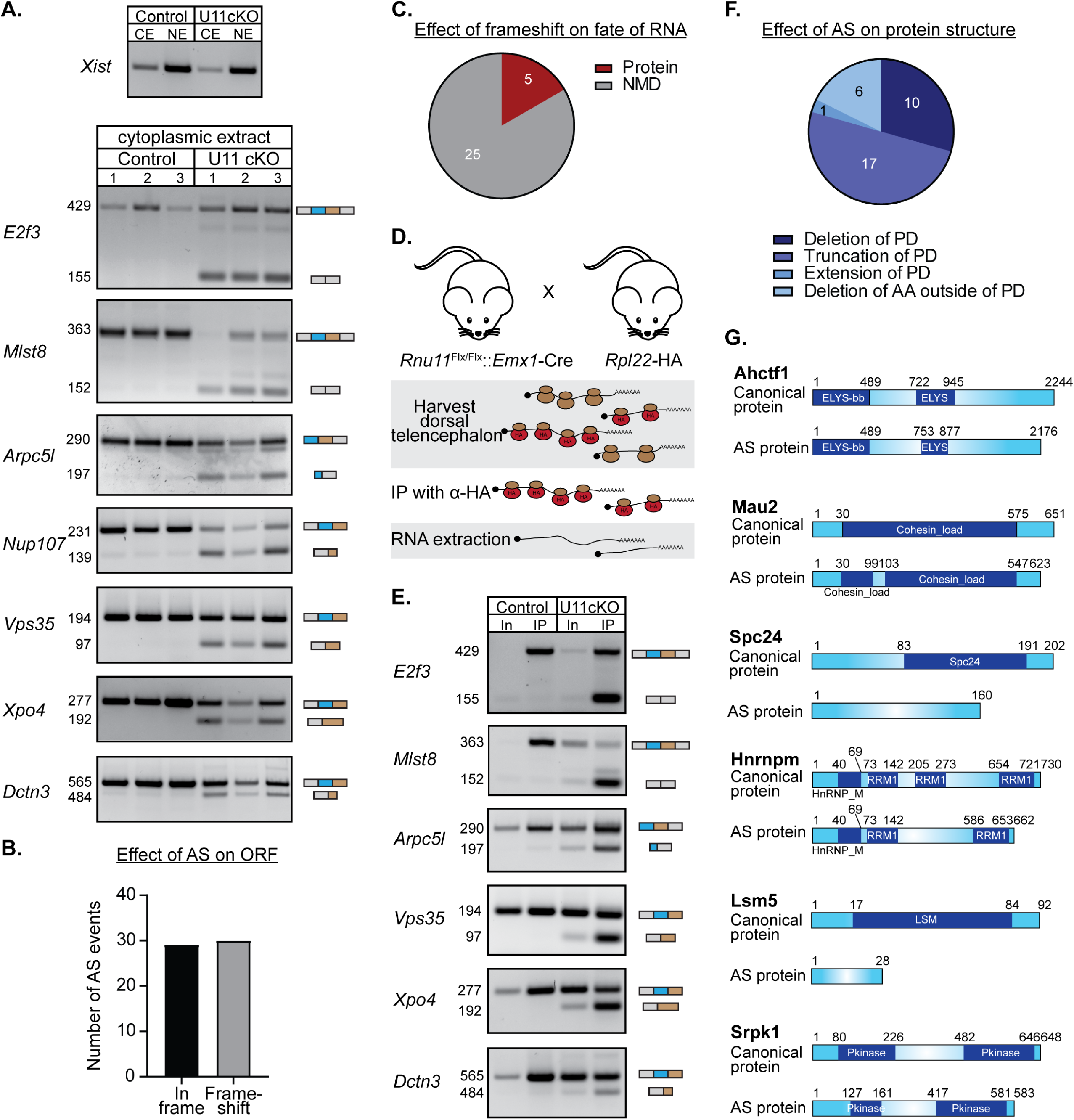
Alternatively spliced MIG transcripts are exported to the cytoplasm and bound to polysomes. **(A)** Agarose gel images showing the nuclear lncRNA *Xist* as a fractionation control; shown below are alternatively spliced RT-PCR products in the cytoplasmic fraction of the E12 dorsal telencephalon **(B)** Bargraph with the predicted effects of upregulated AS around minor introns in the U11 cKO on the ORF. **(C)** Piechart with the number of AS events resulting in a premature stop codon that are predicted to trigger NMD. **(D)** Schematic of experimental design to isolate RNA bound to polysomes in the U11 cKO Ribotag mouse. **(E)** Agarose gel images of alternatively spliced RT-PCR products in whole extract (In, Input) and bound to polysomes (IP) of E14 control and U11 cKO dorsal telencephalon. **(F)** Piechart with the effect of AS on protein domains of affected MIGs. PD=protein domain; AA=amino acid **(G)** Schematics showing altered protein domains due to AS around minor introns. Numbers above indicate amino acids that encode for each protein domain.

Next, to explore whether a subset of the alternatively spliced MIG transcripts was indeed translated, we crossed our U11 cKO mice to Ribotag mice (Sanz et al, 2009). RT-PCR analysis on RNA extracted from immunoprecipitated (IP) polysomes showed the presence of all interrogated alternatively spliced MIG transcripts in the E14 U11 cKO embryos (Fig. 4D-E). Even MIG transcripts that were predicted to undergo NMD by bioinformatics analyses were bound to polysomes (Fig. 4E). Thus, we hypothesized that the alternatively spliced MIG transcripts might produce novel protein isoforms. Therefore, we next determined the effect of the AS on protein structure, by translating the ORF *in silico*, which revealed that the majority of AS events would result in a truncation of the protein (Fig. 4F-G). Notably, 50% of the AS events would result in a truncation of protein domains, whereas 29% of the AS events would result in the removal of one or more entire protein domains (Fig. 4F-G). Thus, we hypothesized that these alternatively spliced protein products of MIGs could contribute to disease pathogenesis.

### Expression of an alternatively spliced MIG transcript in the dorsal telencephalon affects radial glial cell divisions

To gain insight into the biological processes that might be affected by the production of aberrant MIG proteins, we performed functional annotation analysis, which showed a significant enrichment of the GO-term condensed chromosome kinetochore, suggesting that cell cycle may be affected (Fig. 5A). Indeed, *Ahctf1, Spc24, Nup107* and *Dctn3*, the four alternatively spliced MIGs that enriched for this function, play known roles during mitosis (Fig. 4G; 5A-B) (McCleland et al, 2004; Platani et al, 2009; Raaijmakers et al, 2013; Rasala et al, 2006). Previously, we have shown a prolonged pro-metaphase to metaphase transition of U11-null radial glial cells (RGCs) (Baumgartner et al, 2018). Moreover, we found that loss of U11 predominately affected the survival of self-amplifying RGCs. These phenotypes could either be the result of the loss of functional MIG proteins, or of a toxic gain of function acquired by aberrant MIG isoforms. To test whether AS of MIG transcripts contributed to the production of proteins with a toxic gain-of-function, we employed *in utero* electroporation (IUE) to express the full-length alternatively spliced *Dctn3* transcript (Dctn3-AS-Myc) and GFP in E14.5 wild-type embryos (Fig. 5C; S5A). At this timepoint, the dorsal telencephalon predominantly consists of neurons in the cortical plate and intermediate zone, but also contains two types of neural progenitor cells: RGCs and intermediate progenitor cells (IPCs) (Gotz & Huttner, 2005). While IPCs are mostly found in the sub-ventricular zone (SVZ), RGCs are located in the ventricular zone (VZ) and consequently take up the electroporated plasmids (Camarero et al, 2006). We found that overexpression of Dctn3-AS-Myc resulted in a significantly larger percentage of GFP^+^ cells in the VZ/SVZ, as well as significantly fewer GFP^+^ cells in the IZ (Fig. 5D). To determine the identity of these cells, we performed immunofluorescence with antibodies against Pax6 and Tbr2, to mark RGCs and IPCs, respectively. We found that overexpression of Dctn3-AS-Myc resulted in a significantly higher number of Pax6^+^/GFP^+^ RGCs in the VZ/SVZ as a percentage of all GFP^+^ cells in these layers compared to the control (Fig. 5E). In contrast, there was no change in the number of Tbr2^+^/GFP^+^ IPCs in the VZ/SVZ between control and Dctn3-AS-Myc (Fig. S5B-C). Thus, this data suggests that aberrant expression of the alternatively spliced transcript of *Dctn3* is sufficient to alter the types of cell divisions made by RGCs.

**Figure 5.**
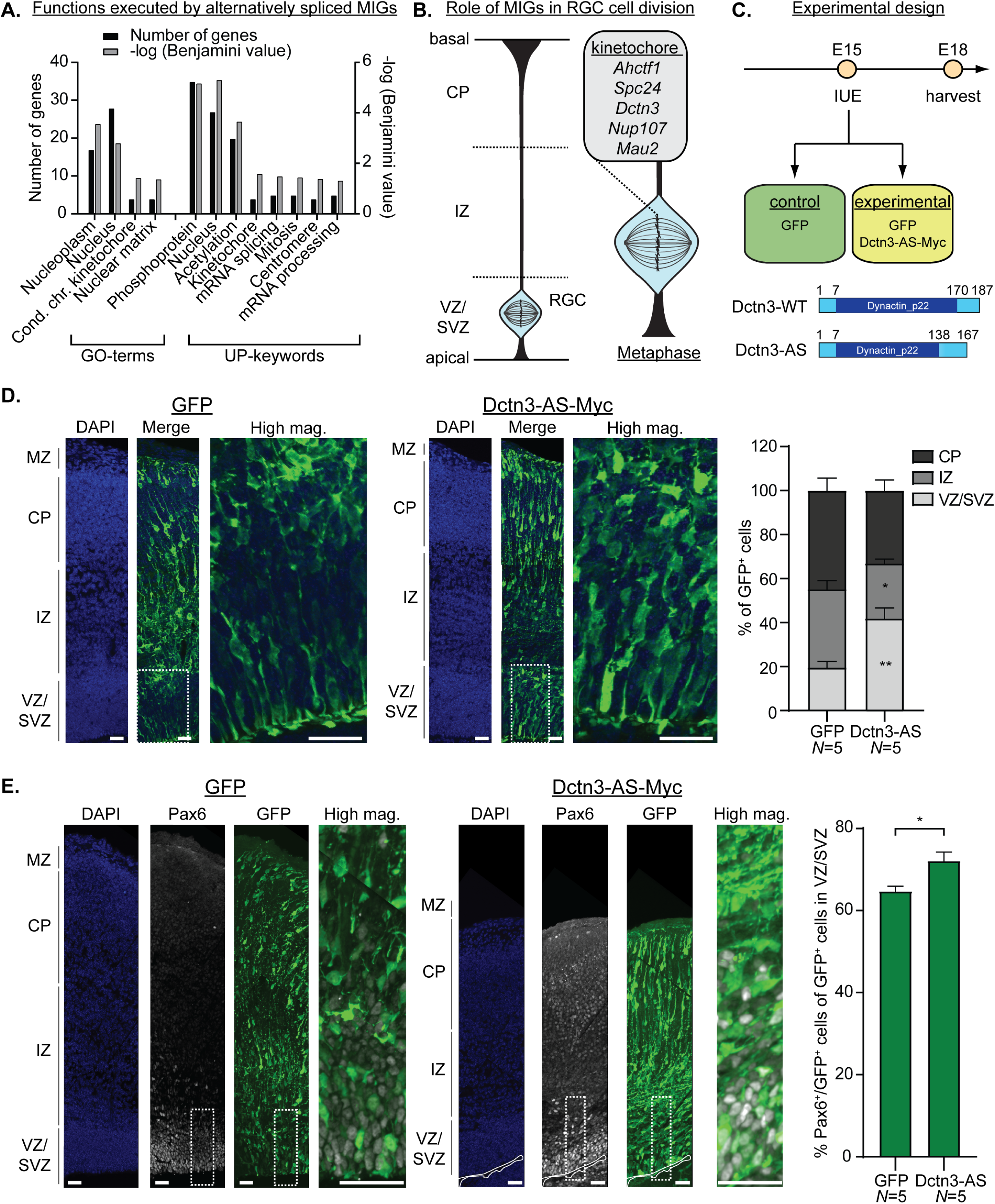
Expression of alternatively spliced MIG transcripts in the dorsal telencephalon affects radial glial cell divisions. **(A)** Bargraph with functional annotation of MIGs with elevated AS around minor introns in the U11 cKO. **(B)** Schematic of layers in developing cortex and the location of radial glial cells (RGCs). On the right is a schematic of an RGC in metaphase and the MIGs enriching for the kinetochore function in **(A). (C)** Experimental design for *in utero* electroporations (IUE). Shown below is a schematic of the protein domains of the two isoforms of Dctn3. **(D-E)** Immunofluorescence for GFP (**D**) and Pax6 (**E**) on E17.5 cortices electroporated with GFP and Dctn3-AS-Myc. Higher magnification image of the insets is shown next to each panel, with the quantification on the right. DAPI marks all nuclei. Scale bars=30um. Statistical significance was determined by two-tailed Student’s t-test. **\*=***P*<0.05; **\*\*=***P*<0.01. CP=cortical plate; IZ=intermediate zone; VZ=ventricular zone; SVZ=subventricular zone. See also Figure S5.

### U11-59K (PDCD7) mediates exon-definition interactions between the major and minor spliceosome

Since these aberrant alternatively spliced transcripts might contribute to disease pathogenesis, we wanted to understand how AS around minor introns was regulated. Given that minor spliceosome inhibition led to an increase in AS around minor introns, we predicted that the AS was executed by the major spliceosome. Indeed, analysis of the consensus sequences resulting from the newly used splice sites in the U11 cKO showed known major-type consensus sequences at the 5’SS and 3’SS, suggesting that the major spliceosome executes AS around minor introns when the minor spliceosome is disrupted (Fig. S6A). This suggests that normally, the minor spliceosome promotes the splicing of the upstream major intron by the major spliceosome, and that its disruption results in exon skipping. As such, these findings reveal crosstalk between the minor and major spliceosome. Therefore, we next sought to identify splicing and/or AS factors that might be involved in this crosstalk. For this, we designed an AS mini-gene construct with a minor intron and the flanking major introns of the MIG *Mlst8*, such that disruption of the interactions between major and minor spliceosome would result in the elevated production of an alternatively spliced transcript consisting of exons 1 and 4 (Fig. 6A’; S6B). We then leveraged this readout to perform a siRNA screen against 55 known splicing and AS factors, followed by RT-PCR analysis for the Mlst8 construct. Quantification of the alternatively spliced product as a ratio of the canonically spliced transcript (%MSI_AS_), showed the largest increase resulted from siRNAs against *PDCD7* (U11-59K), *RNPC3* (U11/U12-65K) and *ZRSR2* (Urp) (Fig. 6A; S6C). After successful confirmation of downregulation of these genes, we repeated the screen with siRNAs against *PDCD7, RNPC3* and *ZRSR2* in triplicate (Fig. 6B-E). This revealed that AS across the minor intron of *Mlst8* was indeed significantly >2-fold upregulated upon inhibition of these splicing factors, compared to the scrambled siRNA (Fig. 6D-E). Thus, the crosstalk between the major and minor spliceosome might not be through the investigated AS factors, but directly through protein components of the minor spliceosome.

**Figure 6.**
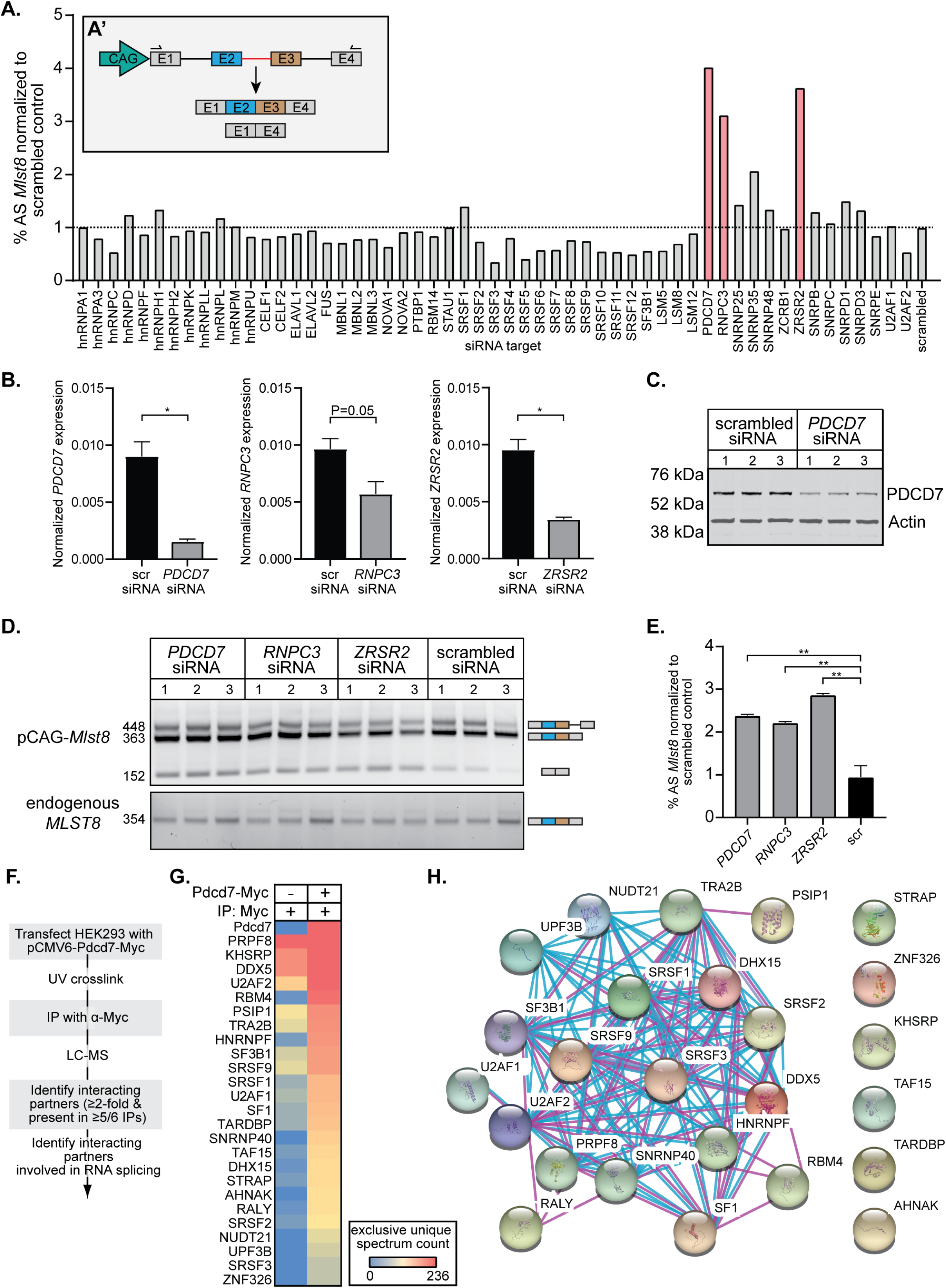
U11-59K (PDCD7) mediates exon-definition interactions between the major and minor spliceosome. **(A)** Bargraph showing the %MSI_AS_ of the *Mlst8* splicing reporter, in response to siRNA against splicing factors. Inset shows schematic of the *Mlst8* splicing reporter and the resulting products. **(B)** qRT-PCR results with expression levels of *PDCD7, RNPC3* and *ZRSR2*, normalized to *GAPDH*. Significance was determined by Student’s t-test. **(C)** Immunoblot for PDCD7 and Actin. **(D-E)** Agarose gel image of the *Mlst8* splicing reporter RT-PCR products resulting from downregulation of minor spliceosome proteins **(D)** and quantification of the %MSI_AS_ using ImageJ **(E).** Significance was determined using one-way ANOVA, followed by post-hoc Tukey test. **(F)** Schematic of experimental design for the transfection and IP of Pdcd7. **(G)** Heatmap of the exclusive unique spectrum count of the Pdcd7-interacting proteins involved in splicing. **(H)** STRING-network of known interactions between Pdcd7-interacting proteins involved in splicing. **\*=***P*<0.05; **\*\*=***P*<0.01. See also Figure S6.

Crosstalk between the major and minor spliceosome has been suggested before to explain the exon-definition model in MIGs (Wu & Krainer, 1996). According to the exon-definition model, the 3’ end of an intron is identified through exon-bridging interactions between the U2AF complex, which binds to the polypyrimidine tract (PPT) of major introns, and U1 snRNP, which binds to the 5’SS of a downstream major intron (Hoffman & Grabowski, 1992; Robberson et al, 1990). For genes consisting exclusively of major introns, it is known that U1-70K is crucial for the maintenance of the exon-bridging interactions (Jamison et al, 1995). However, since the minor spliceosome does not contain U1-70K, nor the U2AF complex, it remained unclear how exon-definition complexes could be formed between major and minor introns in MIGs (Will et al, 1999). Since PDCD7 (U11-59K) is the only U11 snRNP component that showed increased AS (Fig. 6E), we hypothesized that it could perform an analogous role to U1-70K in the maintenance of exon-definition interactions involving the minor spliceosome. To test this hypothesis, we sought to identify Pdcd7-interacting proteins by transfecting HEK293 cells with pCMV-Pdcd7-Myc, followed by immunoprecipitation and mass spectrometry (Fig. 6F). To determine the most robust interacting partners, we curated proteins with ≥2-fold higher spectrum counts in the Pdcd7-Myc condition compared to a negative control, which resulted in a list of 175 proteins (Fig. 6F; S6D-E). Since we wanted to study exon-bridging interactions, we then focused on proteins with a known role in splicing (Fig. 6F). This resulted in a list of 25 proteins, of which many are known to interact with each other (Fig. 6G-H). Specifically, we detected several proteins that have been identified in the major spliceosome E complex, such as SF1, SRSF1, SRSF2, SRSF3, SF3B1, U2AF1 and U2AF2 (Fig. 6H) (Makarov et al, 2012). Moreover, the U5 snRNP components SNRNP40 and PRPF8 co-immunoprecipitated with Pdcd7. In all, these data suggest that U11 snRNP normally interacts with the major spliceosome E complex through PDCD7. Thus, when the minor spliceosome is inhibited these exon-bridging interactions are disrupted, which then results in the AS around minor introns that is executed by the major spliceosome.

## Discussion

### Relationship between AS around minor introns and severity of symptoms in minor spliceosome-related diseases

While only responsible for the splicing of 0.5% of all introns, the importance of the minor spliceosome in development is underscored by the diseases MOPD1, Roifman syndrome, Lowry-Wood syndrome, early-onset cerebellar ataxia and IGHD (Argente et al, 2014; Edery et al, 2011; Elsaid et al, 2017; Farach et al, 2018; He et al, 2011; Merico et al, 2015). The underlying molecular etiology in all these diseases is inhibition of the minor spliceosome, even though the disease-causing mutations are found in different minor spliceosome components. Consequently, these diseases can be characterized by several overlapping symptoms such as microcephaly, developmental delays and growth retardation (Olthof et al., *in press* elsewhere). In the *RNU4ATAC*-related diseases these symptoms are found on a spectrum of severity, where individuals with MOPD1 are most severely affected, and individuals with Lowry-Wood syndrome the least (Shelihan et al, 2018). This suggests the presence of genotype-phenotype relationships that might be informed by the effect of the specific mutations on the secondary structure of U4atac snRNA and therefore inhibition of the minor spliceosome. As a result, differences in the level and types of minor intron mis-splicing might contribute to the differences in phenotype severity. Previous transcriptomic analyses of PBMCs from individuals with Roifman syndrome and MOPD1 have revealed widespread minor intron retention in a shared subset of MIGs, but the authors also noted these samples were hard to compare due to differences in age, cell type, sequencing depth and sex (Cologne et al, 2019). Fortuitously, the RNAseq we performed was on the PBMCs from the individuals with Roifman syndrome and Lowry-Wood syndrome, who were both males, with a similar sequencing depth. This allowed us to compare the effect of the different *RNU4ATAC* mutations on the retention and AS of minor introns. Specifically, the individual with Roifman syndrome described in this manuscript contains one variant in the stem II loop of U4atac, which is important for base-pairing with U6atac snRNA, while the other variant is located in the Sm binding domain of U4atac snRNA (Fig. S1A). In contrast, both mutations in the individual with Lowry-Wood syndrome are located in or near the Sm binding domain of U4atac (Fig. S1A). Our analysis revealed a significant overlap in the number of MIGs that showed minor intron retention in both Roifman syndrome and Lowry-Wood syndrome (Fig. 1D). However, the number of retained minor introns was higher in the clinically more severe Roifman syndrome compared to Lowry-Wood syndrome (Fig. 1A, 1D). These findings support the notion of genotype-phenotype relationships that are informed by the level of minor intron mis-splicing. Importantly, our analysis included heterozygous carriers for Roifman syndrome and Lowry-Wood syndrome, which showed that one mutant *RNU4ATAC* allele is not sufficient to result in aberrant minor intron splicing (Fig. 1A-B).

In addition to minor intron retention, we observed a large number of AS events in the individual with Roifman syndrome, but not in the individual with Lowry-Wood syndrome (Fig. 2A). This suggests that the manner in which U4atac snRNA, and in turn the minor spliceosome, is disrupted, might inform whether minor introns are retained and/or alternatively spliced. Specifically, our results suggest that disruption of the Sm binding domain in U4atac, which might reduce the levels of mature U4atac snRNP but not affect the base-pairing with U6atac snRNA, would inhibit the minor spliceosome such that it results in minor intron retention (Shukla et al, 2002). In contrast, mutations in stem II loop of U4atac snRNA, which is important for base-pairing with U6atac snRNA, inhibits the minor spliceosome such that AS around minor introns was elevated (Shukla et al, 2002). Thus, the maintenance of base-pairing interactions between U4atac and U6atac snRNA might be necessary to inhibit AS around minor introns. Together, the elevated levels of AS around minor introns in Roifman syndrome compared to Lowry-Wood syndrome, led us to propose that alternatively spliced MIG isoforms may contribute to the severity of symptoms such as microcephaly in Roifman syndrome (Fig. 2A, S2A).

The hypothesis that the mode of minor spliceosome inhibition informs the levels and types (retention vs. AS) of minor intron mis-splicing is further reinforced by data from individuals with early-onset cerebellar ataxia. These individuals harbor mutations in *RNU12*, which led to elevated AS around minor introns, as reflected by RT-PCR analysis (Fig. 2D). The upregulation of exon skipping, which was the most upregulated AS category in individuals with minor spliceosome-related diseases, was surprising, as previous reports did not find any elevated exon skipping in individuals with Roifman syndrome and MOPD1 (Fig. S2B) (Cologne et al, 2019; Merico et al, 2015). Given that most AS events around minor introns are solely detected upon minor spliceosome inhibition, they may not have been identified previously and may therefore have been missed by bioinformatics pipelines that focus on annotated AS events. Indeed, using our custom bioinformatics pipeline to detect *de novo* AS events around minor introns, we observed that inhibition of all minor spliceosome snRNAs resulted in elevated AS (Fig. 2A, 3A, 3D).

### Alternatively spliced MIG isoforms are not degraded and might be toxic

While our studies begin to elucidate the effect of minor spliceosome inhibition on the splicing and AS of MIGs, it remained unclear how the expression of these aberrant isoforms would impact disease pathogenesis. Bioinformatics analysis showed that many of the AS events observed in the U11 cKO were predicted to result in a premature stop codon, which was confirmed by Sanger sequencing (Fig. 3E, S4B, 4B-C). Unexpectedly, these alternatively spliced MIG transcripts escaped both nuclear degradation as well as NMD, and were instead bound to polysomes (Fig. 4A, 4E). While it is unclear how these transcripts escaped surveillance, we postulate that minor spliceosome inhibition alters the splicing of MIGs such as *Upf1, Ncbp1, Ncbp2, Ice1*, which are crucial components of the NMD pathway (Fig. 2C-D, 3B-C, S4A; Dataset 1-6) (Baird et al, 2018; Baumgartner et al, 2018; Isken & Maquat, 2008). Moreover, several MIGs are involved in nuclear degradation pathways, such as *Exosc1, Exosc2, Exosc5*, and *Exosc9* (Zinder & Lima, 2017). This might explain why we, and others, have shown the presence of aberrant MIG transcripts without their concomitant downregulation (Fig. 1D-G, 2C-D, 4A) (Baumgartner et al, 2018; Markmiller et al, 2014; Merico et al, 2015).

The presence of MIG transcripts in polysomes suggested that they are translated, thereby resulting in the production of aberrant proteins (Fig. 4E). These can either be non-functional and lead to proteasomal degradation, and/or be toxic and result in a gain-of-function. Evidence for the latter possibility comes from our experiments where we expressed the alternatively spliced transcript of Dctn3 in the developing cortex of wild-type embryos (Fig., 5C-E). Dctn3 has been shown to play a crucial role in mitosis and here we have shown that overexpression of the alternatively spliced isoform of Dctn3 resulted in an increased number of Pax6^+^/GFP^+^ cells of all GFP^+^ cells in the VZ/SVZ (Fig. 5E) (Raaijmakers et al, 2013). These results suggest that expression of the alternatively spliced Dctn3 isoform altered radial glia cell divisions, such that RGCs undergo more self-amplifying proliferative divisions or fewer differentiative divisions (Fig. 5D-E). This proof of principle experiment shows the potential toxicity of the expression of a single aberrant MIG isoform. However, since the number of expressed aberrant MIG isoforms is much higher in both individuals with minor spliceosome-related diseases and our U11 cKO mouse, their phenotype might be further exacerbated compared to the results of our IUE experiments. In all, these findings suggest that AS around minor introns can result in the production of toxic proteins, which might contribute to the microcephaly phenotype observed in the U11 cKO mouse and individuals with minor spliceosome-related diseases (Baumgartner et al, 2018). Specifically, these findings shed light on a potential mechanism in which the severity of microcephaly observed in individuals with Roifman syndrome is linked to increased expression of aberrant MIG isoforms compared to individuals with Lowry-Wood syndrome (Fig. 2A, S2A).

### Exon-bridging interactions between the major and minor spliceosome

The AS around minor introns we observed when the minor spliceosome was inhibited, was executed by the major spliceosome, suggesting cross-talk between these two spliceosomes (Fig. S6A). Indeed, interactions between the major and minor spliceosome were first proposed in 1996, when Wu and Krainer discovered that binding of U1 snRNP to a downstream major-type 5’SS would enhance the splicing of an upstream minor intron in the *SCN4A* splicing construct (Wu & Krainer, 1996). This is consistent with the idea that splicing in vertebrates occurs through exon-definition interactions, whereby U1 snRNP bound to a 5’SS can interact with SR proteins and the U2AF complex that is bound to the upstream PPT/3’SS (Fig. 7) (Hoffman & Grabowski, 1992; Robberson et al, 1990; Wu & Maniatis, 1993). This exon-bridging interaction promotes the recruitment of U2 snRNP to the correct BPS and therefore enhances the splicing of the upstream intron (Fig. 7) (Ruskin et al, 1988). While this model has been used to describe major intron splicing, data from Krainer suggested that this model would also extend to minor intron splicing, which was later confirmed in plants (Lewandowska et al, 2004; Robberson et al, 1990; Wu & Krainer, 1996). However, it remained unclear whether binding of U11/U12 di-snRNP to minor introns would act analogous to U1 snRNP, and enhance the splicing of the upstream major intron. We found that loss of U11 snRNA did indeed result in the upregulation of exon skipping, suggesting that lack of U11 binding to the 5’SS of minor introns, resulted in failure to use the 3’SS of upstream major introns by the major spliceosome (Fig. 3D-F). Thus, these data suggest that U11 snRNP plays a role in mediating exon-definition interactions for a subset of MIGs. Similarly, binding of U11 snRNP to U11 snRNP splicing enhancer (USSE) sequences in introns of *SNRNP48* and *RNPC3* has been shown to activate the usage of an upstream major-type 3’SS (Verbeeren et al, 2010). Thus, the upregulation of AS around minor introns in response to U11 loss can easily be reconciled with an analogous function of U11 snRNP to U1 snRNP in maintaining exon-definition interactions (Fig. 3D-F). However, AS around minor introns was also elevated in response to inhibition of the minor spliceosome when U12, U4atac, and U6atac snRNAs were disrupted (Fig. 3A-C). Thus, the successful assembly and activity of the minor spliceosome as a whole might also regulate the major spliceosome.

**Figure 7.**
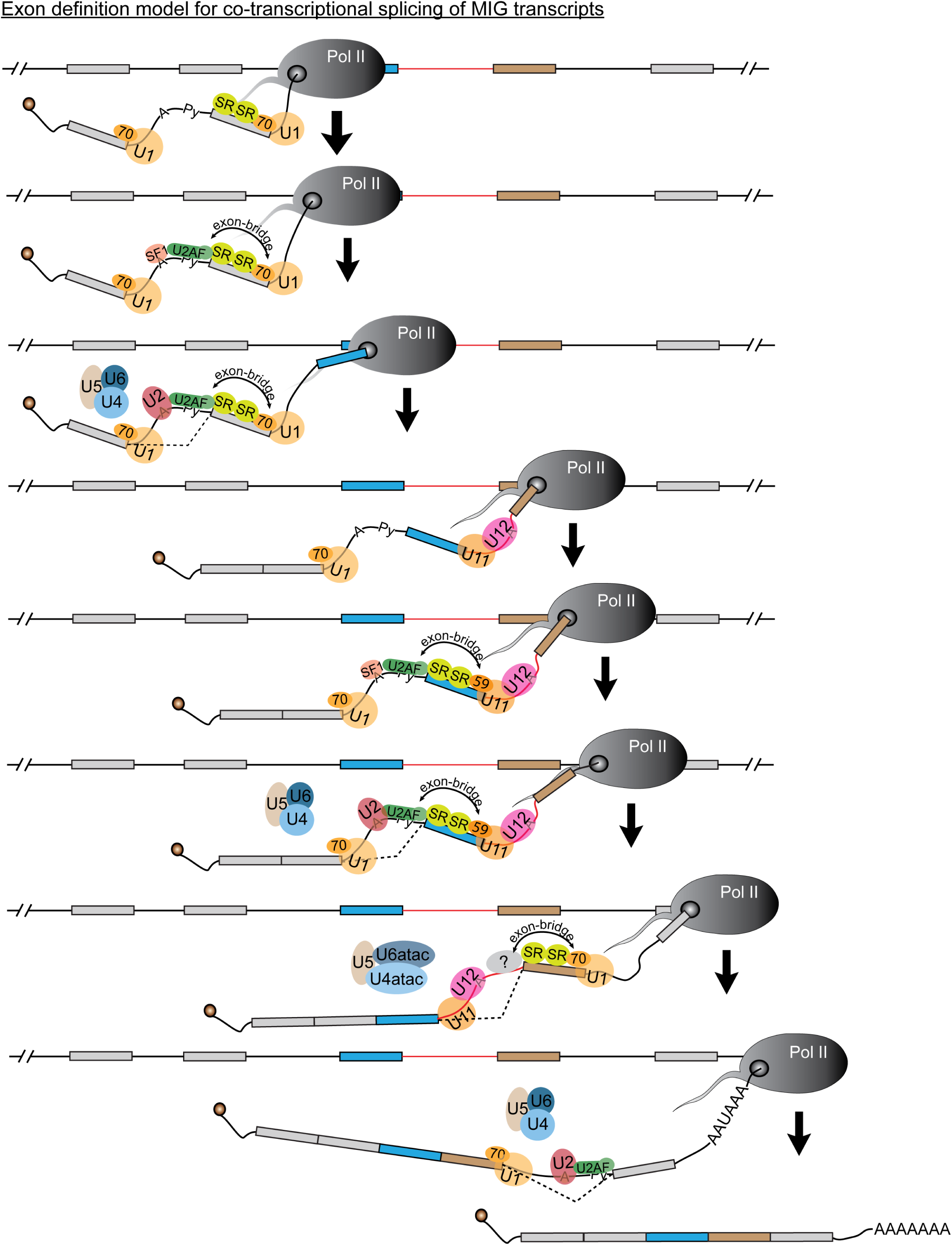
Exon-definition model for splicing of introns in MIGs. Schematic depicting the co-transcriptional splicing of major and minor introns in MIG transcripts. Minor intron is shown as red line, major introns are shown as black lines.

### U11-59K interacts with the major spliceosome

The exon-bridging interactions between major spliceosomes uses U1-70K, which suggests that similar splicing and/or AS factors might play a role to establish exon-definition interactions between the minor and the major spliceosome (Jamison et al, 1995). Based on previous reports that *SNRNP35* (U11-35K) might be the functional analog of U1-70K, we expected that its downregulation by siRNA would result in elevated AS of the *Mlst8* minigene construct (Lorkovic et al, 2004; Will et al, 2004). Indeed, AS levels were upregulated >2-fold upon knockdown of snRNP35, but we saw the largest increase in AS upon downregulation of *PDCD7* (U11-59K), *RNPC3* (U11/U12-65K) and *ZRSR2* (Urp) (Fig. 6A). ZRSR2 is a component of both the major and minor spliceosome, and is thought to play a role in 3’SS selection of minor introns (Shen et al, 2010; Will et al, 1999). In contrast, both PDCD7 and RNPC3 are part of the seven unique minor spliceosome proteins (Will et al, 2004). Since PDCD7 is part of the U11 snRNP and has a highly specific antibody, which gave us confidence in the efficacy of the siRNA, we focused on this protein (Fig. 6C). Immunoprecipitation of PDCD7 followed by LC-MS/MS resulted in the detection of many interacting proteins that included both RNA splicing factors and non-splicing factors (Fig. 6G, S6E). This finding suggests that PDCD7 plays additional roles besides minor intron splicing, which is also evidenced by findings that PDCD7 transactivates E-cadherin expression and is involved in apoptosis of T-cells (Park et al, 1999; Peng et al, 2018). Our focus here was the role of PDCD7 as part of the minor spliceosome, and therefore we curated those interacting proteins known to play a role in RNA splicing. This revealed that PDCD7 interacts with proteins of the SR-protein family, but also SF1, SF3B1, U2AF1 and U2AF2 (Fig. 6G-H, 7). These proteins have all been shown to be crucial components of the exon-bridging complex between major introns (De Conti et al, 2013). Here, we show that they also participate in exon-definition interactions between the minor and major spliceosome. (Fig. 7) Together, these findings for the first time reveal a potential reciprocal regulation of the two spliceosomes to regulate MIG expression.

Overall, our work is revealing the complex regulation of splicing and AS of MIGs through coordinated action of the major and the minor spliceosome, which is in line with the exon-definition model. Given that minor introns and the minor spliceosome evolved very soon after major introns, and are highly conserved across species, the regulated cross-talk is consistent with the idea of co-evolution of the exon-definition interactions (Russell et al, 2006). Inherent to these interactions is the means to regulate the splicing and AS of many MIGs that are essential for survival, cell cycle and other functions (Baumgartner et al, 2019). As the involvement of the minor spliceosome in diseases such as amyotrophic lateral sclerosis, myelodysplastic syndrome, and others are being discovered, understanding the regulation of MIG expression will prove to be invaluable (Fertig et al, 2009; Madan et al, 2015; Reber et al, 2016).

## Materials and methods

### Animal husbandry

All mouse procedures were performed according to the protocols approved by the University of Connecticut Institutional Animal Care and Use Committee (IACUC), which ensures adherence to the U.S. Public Health Service Policy on the treatment of laboratory animals. The generation of the U11 cKO mouse has been described previously (Baumgartner et al, 2018). To ablate U11 snRNA in the developing cortex, we crossed the *Rnu11* cKO mouse to *Emx1-Cre*^*+*^ mice (Gorski et al, 2002). These crosses resulted in E12 control embryos (*Rnu11*^WT/Flx^::*Emx1*-Cre^+/-^) and U11 cKO embryos (*Rnu11*^Flx/Flx^::*Emx1*-Cre^+/-^). To isolate RNA bound to polysomes, we crossed our U11 cKO mice with *Rpl22*-HA^+^ mice, and harvested E14 embryos (Sanz et al, 2009). IUE procedures were performed on E14.5 timed pregnant CD1 female mice obtained from Charles River.

### Human subjects

Informed consent was obtained from individuals with mutations in *RNU4ATAC (N*=2), their unaffected carrier parents (*N=*3), and unrelated healthy controls (*N*=3) using a protocol approved by the Institutional Review Board committee at the CHU Sainte-Justine. The phenotypic description of the individual with Lowry-Wood syndrome described in this manuscript had previously been published (Magnani et al, 2009; Shelihan et al, 2018). In contrast, the individual with Roifman syndrome described in this manuscript had not previously been reported. This individual experienced asymmetric intra-uterine growth retardation during pregnancy along with other ultrasound anomalies (including a suspected aortic coarctation, which was later not confirmed). Prenatal investigations revealed a normal array CGH (Comparative Genome Hybridization) and a customized Noonan syndrome panel revealed an inherited *KMT2D* variant of uncertain significance. Given that this variant was inherited, it was deemed non-pathogenic. The patient was born at term with a weight (2320g) and length (42cm) both below the third percentile. Moreover, the head circumference was at the 50^th^ percentile. He had micromelia and brachydactyly, and a skeletal survey showed platyspondyly, irregular metaphyses and delayed epiphyseal ossification (i.e. a clinical presentation of spondyloepimetaphyseal dysplasia (SEMD)). A SEMD panel then revealed two *RNU4ATAC* mutations *in trans* (Fig. S1A), which have previously been reported in another individual with Roifman syndrome (Dinur Schejtur, 2016). The patient was then evaluated by additional clinicians who determined the presence of low titers of antibodies against tetanus following immunization. Therefore, he is now, at nine months of age, supplemented with immunoglobins. He has not had a serious infection to date and is doing well.

The individuals with mutation in *RNU12* (*N*=5) and their unaffected carrier parents (*N*=3) mentioned in this manuscript have been described previously (Elsaid et al, 2017). Informed consent from these families was obtained using protocols approved by Institutional Review Board committee at Weill Cornell Medicine-Qatar and Hamad Medical Corporation.

### Cell culture

HEK293T cells were cultured in Dulbecco’s modified Eagle’s medium (DMEM), supplemented with 10% fetal bovine serum (FBS), 5% sodium pyruvate and penicillin (100 IU/mL) and streptomycin (100ug/mL). Cells were maintained at 37 °C and 5% CO_2_.

### In utero electroporation (IUE) and immunofluorescence

In utero electroporation was performed as described previously (Chen et al, 2014). Briefly, timed pregnant (E14.5) CD1 females were anesthetized by intraperitoneal (i.p.) injection of a mixture of 90mg/kg bodyweight ketamine and 10mg/kg xylazine. After exposure of the uterine horns, ∼1 ul of plasmid DNA (1.5ug/ul) mixed with Fast Green was injected into the lateral ventricles of the embryos using a glass micropipette. For the control condition this included a pCAG-GFP plasmid, whereas the experimental condition consisted of an equal mixture pCAG-GFP and pCAG-Dctn3AS-Myc. Using a square-pulse electroporator and electrode paddles placed on each side of the embryo’s head, we then applied five 50ms pulses of 35V at 0.1 second intervals. After the electroporations, the uterus was placed back in the abdominal cavity and the wound was surgically sutured. The mice were closely monitored following the surgery and Metacam analgesics was injected subcutaneously once a day for two days. After 72 hours, embryos were harvested and the heads were used to generate 30um coronal cryosections. These were used for immunofluorescence as described in (Karunakaran et al, 2015). Primary antibodies were diluted to 1:300 (chicken anti-GFP; rabbit anti-Tbr2; rabbit anti-Pax6) or 1:500 (mouse anti-Tubb3 and all secondary antibodies).

### Image acquisition and quantification

Processed slides were imaged with a Leica SP2 confocal microscope, where settings for laser intensity, excitation-emission windows, gain and offset conditions were identical between the two conditions on a given slide. Further processing was performed on IMARIS v.8.3.1 (Bitplane) and Adobe Photoshop CS4 (Adobe Systems). All image processing in IMARIS and Photoshop was done using identical settings between the two conditions. Manual quantification was performed using the spot tool in IMARIS and statistical significance was calculated using two-tailed Student’s t-test.

To determine MSI values based on RT-PCR analysis, we employed ImageJ. Band intensity of canonically spliced and alternatively spliced products was calculated and used to determine the MSI value as is done for RNAseq (i.e. intensity AS product / (intensity AS product + intensity canonical product).

### Immunoprecipitation

#### Polysomes

Ribosomal-bound RNA was extracted from the dorsal telencephalons of E14 *Rnu11*^WT/Flx^::*Emx1*-Cre^+/-^::*Rpl22*-HA^+/-^ and *Rnu11*^Flx/Flx^::*Emx1*-Cre^+/-^::*Rpl22*-HA^+/-^ embryos as described previously (Sanz et al, 2009). Briefly, for each litter, the dorsal telencephalons of all embryos of each genotype (∼*N*=4) were pooled and used for dounce homogenization in 1mL polysome buffer (50mM Tris pH7.5, 100mM KCl, 12mM MgCl_2_, 1% Igepal, 1mM DTT, 200U/ml RNAse Inhibitor, 1mg/mL heparin, 100ug/mL cycloheximide, 1X protease inhibitor). 50uL of the dounce homogenized sample was labelled as “input” and added to 350uL RLT buffer with 1% 2-mercaptoethanol (Qiagen), followed by RNA extraction. The remainder sample was centrifuged and the supernatant was extracted. This was then added to HA-coupled protein G Dynabeads (ThermoFisher Scientific, #10004D) and rotated overnight at 4C. Afterwards, beads were washed three times in high salt buffer (50mM Tris pH7.5, 300mM KCl, 12mM MgCl_2_, 1% Igepal, 1mM DTT, 100ug/mL cycloheximide) and 350uL RLT buffer with 1% 2-mercaptoethanol was added to the beads. After vigorous vortexing, the beads were immobilized and supernatant was labelled as “IP”. Fig. 4E contains representative images of results that have been repeated at least four times.

#### Pdcd7-Myc

To identify which proteins Pdcd7 interacts with, we overexpressed a pCMV6-Pdcd7-Myc plasmid (OriGene, #MR214517) in HEK293T cells using GenJet II (SignaGen, #SL100489), as per the manufacturer’s guidelines. As a control, an empty vector was utilized. After 48hrs, cells were washed three times in 1X PBS and crosslinked with UV (365nm) using the Stratalinker 2400. Settings were set to 4000J/cm^2^, followed by 2000J/cm^2^. Cells were dislodged and resuspended in 400ul modified RIPA buffer (50mM Tris-HCl pH8.0, 150mM NaCl, 0.5% SDS and 1X protease inhibitor). The lysate was then pre-cleared for 1 hour by mixing 400uL sample with protein G Dynabeads (ThermoFisher Scientific, #10004D). Afterwards, the pre-cleared lysate was incubated with a primary antibody (goat anti-Myc #ab9132 or goat anti-mouse IgG (H+L) #115-005-003) overnight at 4C. Protein G Dynabeads were then added to the samples, followed by a 3-hour incubation, and washing with 1X PBS. Finally, proteins were extracted by adding 30uL modified RIPA buffer and boiling of the sample at 95C for 3 minutes.

### Mass spectrometry

Protein samples (*N*=6 per condition) were submitted to the Proteomics and Metabolics Core facility at the University of Connecticut for a slightly modified filter-aided sample preparation (FASP) method in a microcon YM-10 10kD molecular weight cutoff (MWCO) filter (Thermo Fisher Scientific) (Wisniewski et al, 2009). Briefly, all IP elutions were diluted to 250 µL final volume using UA buffer (8 M urea in 0.1 M Tris HCl, pH 8.5) and reduced using 25 mM dithiothreitol for 1.5 hr at 37°C. The samples were added to the MWCO filter and spun at 14,000 x g for 40 min, washed with 200 µL UA buffer and spun again using Spin Condition 1 (14,000 x g for 40 min). Cys residues were carbamidomethylated using 50 mM iodoacetamide in UA buffer for 15 min in the dark at 37°C and then centrifuged at Spin Condition A. Proteins were washed using two cycles of resuspension in 100 µL UB buffer (8 M urea in 0.1 M Tris HCl, pH 8.0) and centrifuged using Spin Condition 1 (14,000 x g for 30 min). Proteins were resuspended in 50 µL UB buffer, removed from the MWCO filter and placed into a new 1.7 mL Eppendorf safe-lock tube. The MWCO filter was washed with two aliquots of 50 µL 0.1 M ammonium bicarbonate and pooled to result in a final urea concentration <1 M. The first stage of proteolysis was initiated by adding Endoproteinase LysC (Pierce) at a 1:50 protein:protein (w/w) ratio for 16 hr at 37°C on a thermo shaker (Thermo Scientific). Sequencing Grade Modified Trypsin (Promega) was then added at a 1:50 protein:protein (w/w) ratio and allowed to proceed for an additional 8 hr at 37°C. Enzymatic digestions were quenched using concentrated formic acid to result in a final pH of 2.5 and desalted using C18 Peptide Desalting Spin Columns (Pierce) following manufacturer’s instructions.

Peptide samples were subject to mass analysis using a Thermo Scientific Ultimate 3000 RSLCnano ultra-high performance liquid chromatography (UPLC) system coupled directly to a high resolution Thermo Scientific Q Exactive HF mass spectrometer. An aliquot of each peptide preparation was injected onto a PepMap RSLC C18 analytical column (2 µm, 100 Å, 75 µm x 25 cm, Thermo Scientific) and subject to a 90 min, 300 nL/min reversed-phase UPLC method. Peptides were eluted directly into the Q Exactive HF using positive mode nanoflow electrospray ionization. A data-dependent Top 15 tandem mass spectrometry (MS/MS) acquisition method was used that implemented the following parameters for MS scan acquisition: 60,000 resolution, 1e6 AGC target, maximum ion time of 60 ms, and a 300 to 1800 m/*z* mass range of. MS/MS scan acquisition included the following parameters: 15,000 resolution, maximum ion time of 40 ms, isolation window of 2.0 m/*z*, 30 s dynamic exclusion window, normalized collision energy of 27, a 200 to 2,000 m/*z* scan range, and charge exclusion “on” for all unassigned, +1, and >+8 charged species.

Peptides were identified and quantified via label-free quantification using MaxQuant (v1.6.0.1) and the embedded Andromeda search engine (Cox & Mann, 2008). The raw data was then searched against an in-house generated protein database consisting of the myc-tagged Pdcd7 primary sequence and the entire Uniprot *Homo sapiens* proteome database (identifier UP000005640, accessed 20170422). The following parameters were used for the search: variable modifications oxidation of Met, acetylation of Protein N-termini, and Gln to pyro-Glu conversion, fixed carbamidomethylation of Cys, trypsin enzyme specificity with up to 2 missed cleavages, LFQ quantitation “on”, and a minimum of 5 amino acids per peptide. All results were filtered to a 1% false discovery rate at the peptide and protein levels; all other parameters were kept at default values. MaxQuant-derived output was uploaded into Scaffold Q+S (v 4.0, Proteome Software) for visualization and further analysis.

Only proteins for which exclusive unique spectrum counts were detected in five out of the six Pdcd7 IPs were included for further analysis. Interacting proteins were defined as having a fold change in the sum of exclusive unique spectrum counts between the Pdcd7-Myc and control condition greater than 2. To identify interacting proteins that could play a role in exon-definition, all interacting proteins were submitted to DAVID for functional annotation analysis, and proteins that significantly enriched for the function mRNA splicing were extracted.

### siRNA screen

To knockdown splicing factors in HEK293T cells, we obtained a customized library containing ON-TARGET plus SMARTpool siRNAs from Dharmacon Inc. siRNAs were resuspended in UP H_2_O to a concentration of 2uM. 50pmol of siRNA against each gene, combined with 500ng pCAG-Mlst8 splicing reporter were co-transfected in 24-well plates using DharmaFECT DUO, as per the manufacturer’s guidelines (Dharmacon Inc, #T-2010-01). After 48 hours, cells were harvested in 100uL TRIzol or 100uL modified RIPA buffer (50mM Tris-HCl pH8.0, 150mM NaCl, 0.5% SDS and 1X protease inhibitor). To repeat the experiments with N-value, we obtained siGenome SMARTpool siRNAs for *PDCD7* (#M-012096-00-0005), *RNPC3* (#M-021646-01-0005), and *ZRSR2* (#M-006596-01-0005) and followed the same experimental paradigm as described before.

### Plasmids and splicing constructs

The pCAG-GFP plasmid (Addgene, #11150) was obtained from Addgene and utilized for the design of the pCAG-Mlst8 splicing reporter and the pCAG-Dctn3-AS-Myc plasmid. First, the pCAG-GFP plasmid was cut with *EcoRI* and *NotI* to remove the GFP cassette. The four exons and three introns of Mlst8 were PCR amplified with primers listed in Table S1 and then cloned into the pCAG backbone using the Gibson Assembly Master Mix (NEB, #E2611S). Similarly, the alternatively spliced isoform of Dctn3 was amplified using primers listed in Table S1, followed by cloning into the pCAG backbone using the Gibson Assembly Master Mix (NEB, #E2611S). The pCMV6-Pdcd7-Myc plasmid was obtained from Origene (OriGene, #MR214517).

### Morpholino electroporation

Electroporations were performed as described previously (Matter & Konig, 2005). Briefly, control (5′-CCTCTTACCTCAGTTACAATTTATA-3’) (Younis et al, 2013), U12 (5’-TCGTTATTTTCCTTACTCATAAGTT-3’) (Matter & Konig, 2005), U4atac (5’-CAGGCGTTAGCAGTACTGCCCTCAC-3’), U6atac (5′-AACCTTCTCTCCTTTCATACAACAC-3′) (Younis et al, 2013) and U2 (5’-TGATAAGAACAGATACTACACTTGA-3’) (Matter & Konig, 2005) antisense morpholino oligonucleotide were obtained from GeneTools, Inc and dissolved in water to a final concentration of 0.5 nmol/uL. Approximately 1-2*10^4^ cells were used for each separate electroporation (*N=3* per electroporation condition). Cells were washed with PBS, followed by mixing with 15nmol of morpholino in a 1.5mL Eppendorf tube. After a 10min incubation at room temperature, cells were transferred to a 4-mm-gap electroporation cuvette and electroporated using a Bio-Rad Gene Pulser at 200V for 50ms. Cells were then plated in 10mL growth medium for 6 hours. Afterwards, cells were washed once in PBS and plated in new growth medium supplemented with 200uM 5’-ethynyl uridine (EU) (ThermoFisher Scientific, #C10365). Two hours later cells were harvested and homogenized in 1 mL TRIzol (ThermoFisher Scientific, #15596018), followed by extraction of total RNA. For isolation of nascently transcribed, EU-pulsed RNA, 5ug of total RNA was biotinylated using the Click-IT kit (ThermoFisher Scientific, #C10365), per the manufacturer’s instructions. Then, 50ng of biotinylated RNA was pulled down on Dynabeads (ThermoFisher Scientifc, #C10365) and resuspended in 5uL buffer. The nascently transcribed RNA was then immediately used for library preparation as described below.

### RNA extraction, cDNA preparation and PCR analysis

#### U11-null dorsal telencephalon

Dorsal telencephalons were dissected from E12 *Rnu11*^WT/Flx^::*Emx1-Cre*^+/-^ (*N=3*) and *Rnu11*^Flx/Flx^::*Emx1-Cre*^+/-^ (*N=3*) embryos and individually used for RNA extraction. The tissue was triturated in 100uL TRIzol (ThermoFisher Scientific, #15596018) and RNA was extracted using the DirectZOL RNA Miniprep Plus Kit (Zymo Research, #R2072), per the manufacturer’s instructions. 500ng of total RNA was then used for cDNA synthesis, as described previously (Kanadia et al, 2006) and 25ng of cDNA was used for reverse transcriptase PCR (RT-PCR) analyses.

#### HEK293T cells

For the experiments described in Figure 2, HEK293T cells were resuspended in 1mL TRIzol (ThermoFisher Scientific, #15596018) and RNA was extracted using phenol:chloroform, as per manufacturer’s instructions. For RT-PCR analyses, 1ug of total RNA was used for cDNA synthesis. For the siRNA screen, HEK293T cells were resuspended in 100uL TRIzol and RNA was extracted using the DirectZOL RNA Microprep kit (Zymo Research, #R2062). 100ng of total RNA was then used for cDNA synthesis and 25ng of cDNA was used for RT-PCR analysis. To confirm downregulation of *PDCD7, RNPC3* and *ZRSR2*, we performed quantitative RT-PCR (qRT-PCR) analysis on 25ng cDNA. The Cq values were then normalized to the expression of *GAPDH*.

#### PBMCs from individuals

PBMCs from individuals were pelleted and washed three times with 1X PBS. Afterwards, the pellet was resuspended in 100uL TRIzol and the RNA was extracted using the DirectZOL RNA Microprep kit (Zymo Research, #R2062). 100ng of total RNA was then used for cDNA synthesis and 20ng of cDNA was used for RT-PCR analysis.

### Bioinformatics analysis

#### Library preparation

Total RNA was depleted from rRNA using the RiboZero kit (#MRZH116) by the Center for Genome Innovation at the University of Connecticut. A cDNA library was then prepared using the Illumina TruSeq Stranded Total RNA Library Sample Prep Kit (#RS-122-2201) and sequenced on the Illumina NextSeq 500. This resulted in 60-100 million paired-end 151 bp reads per sample.

#### Gene expression analysis

Reads from each sample were aligned to the hg38 genome (UCSC genome browser) using Hisat2 as described previously (Baumgartner et al, 2018; Kim et al, 2015). Gene expression was then determined by IsoEM2, followed by differential gene expression by IsoDE2, as described previously (Al Seesi et al, 2014; Baumgartner et al, 2018; Mandric et al, 2017).

#### Intron retention and alternative splicing analysis

Minor intron coordinates were downloaded from the Minor Intron DataBase (MIDB) (Olthof et al, 2019). Coordinates of the flanking major introns were then extracted for the canonical MIG transcripts (as defined by Ensembl). These were then used to determine retention and alternative splicing levels as described previously (Olthof et al, 2019).

#### Functional enrichment analysis

Genes were submitted for functional enrichment analysis to DAVID (Huang da et al, 2009). Only significant GO-Terms (Benjamini-Hochberg adjusted *P*-value<0.05) were reported.

#### Consensus sequence analysis

Novel junction coordinates were extracted for each upregulated AS event in the U11 cKO dorsal telencephalon using BEDTools (Quinlan & Hall, 2010). These were then utilized to extract the novel intronic sequences generated by AS events with the BEDTools getfasta tool. Finally, frequency plots of the annotated and novel splice sites were made using WebLogo (Crooks et al, 2004). If not a single novel exon-exon junction was supported by more than 10 reads, this MIG was excluded from the analysis.

#### ORF analysis

To determine the effect of AS events across minor introns on the ORF, known exon-exon junctions for the canonical MIG transcripts were adjusted to contain the novel junction coordinates using BEDTools (Quinlan & Hall, 2010). These were then used to generate the full coding sequence as well as for *in silico* translation. To predict whether an alternatively spliced transcript with premature stop codon would be targeted for NMD, the location of the stop codon was compared to the annotated last exon-exon junction of each gene. If this was >50nt upstream of the last exon-exon junction, it was predicted to be targeted to NMD, otherwise it was considered to be translated into protein. The effect of AS on protein domains was determined using the pfam database (Finn et al, 2016). Alternatively spliced MIGs that did not have a single novel exon-exon junction supported by more than 10 reads were excluded.

#### Principal component analysis

Principal component analysis was performed based on MIG expression (TPM values), retention levels (%MSI_Ret_) and alternative splicing levels (%MSI_AS_) using the default settings in ClustVis (Metsalu & Vilo, 2015).

#### Quantification and statistical analysis

Statistical details of the experiments can be found in the figures and figure legends, as well as the results section. To determine whether the retention and AS levels of individual introns were significantly different in the U11 cKO, we performed a Student’s t-test. For the morpholino experiments, a one-way ANOVA followed by post-hoc Tukey test was performed. To determine whether major intron retention levels were generally significantly different between control and U11 cKO, we performed a Mann-Whitney U-test. Moreover, statistical differences in expression levels of PDCD7, RNPC3 and ZRSR2 was determined by Student’s t-test. Furthermore, significant differences in cell numbers in the IUE experiments were also determined by Student’s t-test. *P*<0.05 was considered as significant in all analyses.

## Data availability

The datasets generated during this study are available in the following database:

- RNA-seq data: Gene Expression Omnibus GSE96616. (https://www.ncbi.nlm.nih.gov/geo/query/acc.cgi?acc=GSE96616)

## Acknowledgements

We would like to thank Dr. Bo Reese from the Center of Genome Innovation at the University of Connecticut for helping with the RNA sequencing. Moreover, we would like to thank Dr. Jeremy Balsbaugh from the Proteomics and Metabolomics Core at the University of Connecticut for performing the mass spectrometry experiments. Furthermore, we would like to thank previous members of the Kanadia lab, Roy Levit and Shannon Blemings for help with RT-PCR analysis. Moreover, Gabriela Aquino was instrumental for maintenance of the mouse colony. We are also thankful to Jesse Dunnack who trained us to perform the IUE procedures. In addition, we would like to acknowledge Dr. Ion Mandoiu from the Computer Science Engineering Department for creating the necessary infrastructure to perform bioinformatics analysis. Furthermore, we would like to acknowledge Dr. Jean Jacques De Bruycker from the University of Montreal and Dr. Chaim M. Roifman and Dr. Peter Kannu from the University of Toronto as they have been instrumental in the care of the individual with Roifman Syndrome. Finally, we thank Dr. Karen Menuz for insightful comments on the organization of the manuscript. Funding obtained for this study comes from NIH NINDS R01NS102538 and NIH NINDS R21NS096684 to R.N.K.

## Author Contributions

Conceptualization, A.M.O. and R.N.K.; Methodology, A.M.O. and R.N.K.; Software, A.M.O.; Investigation, A.M.O., A.W., M.L., A.C., J.R., P.M.C. and R.N.K.; Resources, A.K.A.A., C.M. and P.C.; Writing – Original Draft, A.M.O. and R.N.K; Writing – Review & Editing, A.M.O., P.M.C. and R.N.K.; Supervision, R.N.K.; Funding Acquisition, R.N.K.

## Supplemental Information

**Figure S1. The effect of *RNU4ATAC* mutations on minor intron retention and MIG expression. (A)** Schematic of the secondary structure of U4atac snRNA and the locations of the mutations in the RS and LWS patient. **(B)** Sashimi plots showing read coverage across minor introns. **(C)** Principal component analysis (PCA) on TPM values of MIGs in patients and controls. Prediction ellipse is drawn such that the probability is 95% that a new observation from the same condition falls inside the ellipse. RS=Roifman syndrome; LWS=Lowry-Wood syndrome.

**Figure S2. Alternative splicing around minor introns is upregulated in Roifman Syndrome. (A)** Venn diagram of the number of AS events that are upregulated in the RS and LWS patient compared to healthy controls and respective carriers. **(B)** Piechart with the distribution of different AS events that are upregulated in the RS and/or LWS patient. **(C)** Sashimi plots showing combined AS usage around minor introns. **(D)** Principal component analysis (PCA) on %MSI_AS_ in patients and controls. Prediction ellipse is drawn such that the probability is 95% that a new observation from the same condition falls inside the ellipse. **(E)** Schematic of the secondary structure of U12 snRNA and the location of the mutations in the early-onset cerebellar ataxia (EOCA) patients. RS=Roifman syndrome; LWS=Lowry-Wood syndrome.

**Figure S3. Inhibition of minor spliceosome snRNAs through morpholinos results in elevated minor intron retention. (A)** Schematic of experimental design for morpholino (MO) electroporations. **(B-E)** Volcano plot showing the delta MSI_Ret_ for retained minor introns upon inhibition of U12 snRNA **(B)**, U4atac snRNA **(C)**, U6atac snRNA **(D)** and U2 snRNA **(E)**. Significance was determined by one-way ANOVA, followed by post-hoc Tukey test. **(F)** Piechart with the distribution of different AS events that are upregulated upon minor spliceosome inhibition.

**Figure S4. Loss of U11 results in elevated exon skipping. (A)** Heatmap of MSI_AS_ values for AS events uniquely detected in the U11 cKO. **(B)** Gel images of RT-PCR products resulting from AS around minor introns. Molecular weight is shown on the left; product schematics are shown on the right. The %MSI_AS_ was calculated using ImageJ.

**Figure S5. Overexpression of alternatively spliced Dctn3 isoform does not affect IPC numbers. (A)** Western blot for Myc on HEK293T cells transfected with pCAG-GFP, pCAG-Dctn3-WT-Myc and pCAG-Dctn3-AS-Myc. **(B-C)** Immunofluorescence for GFP and Tbr2 on E17.5 cortices electroporated with pCAG-GFP (left) or a combination of pCAG-GFP and pCAG-Dctn3-AS-Myc (right) **(B)**, with quantification in **(C)**. DAPI marks all nuclei. Scale bars=30um. Statistical significance was determined by two-tailed Student’s t-test. n.s.=not significant. CP=cortical plate; IZ=intermediate zone; VZ=ventricular zone; SVZ=subventricular zone.

**Figure S6. Identification of proteins interacting with U11-59K. (A)** Frequency logos of the 5’ SS and 3’ SS of alternatively spliced minor introns in the U11 cKO. **(B)** Example of agarose gel image of the *Mlst8* splicing reporter RT-PCR products resulting from downregulation of proteins in the siRNA screen. **(C)** Equation used to determine the normalized %MSI_AS_ in the siRNA screen. **(D)** Western blot for Myc and Pdcd7 on samples transfected with a control or pCMV6-Pdcd7-Myc plasmid and immunoprecipitated with IgG or Myc antibodies. **(E)** Heatmap of exclusive unique spectrum counts of all proteins >2-fold enriched in the Pdcd7-Myc IP samples. Zoomed in subset shown on the right. Proteins involved in splicing are shown in bold.

**Supplemental datasheet 1. Minor intron retention in Roifman and Lowry-Wood syndrome.**

**Supplemental datasheet 2. MIG expression in Roifman and Lowry-Wood syndrome.**

**Supplemental datasheet 3. AS around minor introns in Roifman and Lowry-Wood syndrome.**

**Supplemental datasheet 4. Minor intron retention upon inhibition of minor spliceosome snRNAs by morpholinos.**

**Supplemental datasheet 5. AS around minor introns upon inhibition of minor spliceosome snRNAs by morpholinos.**

**Supplemental datasheet 6. AS around minor introns in U11 cKO mice.**

